# Disease associated protein-protein interaction network reconstruction based on comprehensive influence analysis

**DOI:** 10.1101/2019.12.18.880997

**Authors:** Fei Zhu, Feifei Li, Xinghong Ling, Quan Liu, Bairong Shen

## Abstract

Proteins and their interactions are fundamental to biological system. With the scientific paradigm shifting to systems biology, functional study of proteins from a network viewpoint to get a deep understanding of their roles in human life and diseases being increasingly essential. Although several methods already existed for protein-protein interaction (PPI) network building, the precise reconstruction of disease associated PPI network remains a challenge. In this paper we introduce a novel concept of comprehensive influence of proteins in network, in which direct and indirect connections are adopted for the calculation of influential effects of a protein with different weights. With the optimized weights, we calculate and select the important proteins and their interactions to reconstruct the PPI network for further validation and confirmation. To evaluate the performance of the method, we compared our model with the six existed ones by using five standard data sets. The results indicated that our method outperforms the existed ones. We then applied our model to prostate cancer and Parkinson’s disease to predict novel disease associated proteins for the future experimental validation.

**Author Summary:** The diverse protein-protein interaction networks have dramatic effects on biological system. The disease associated PPI networks are generally reconstructed from experimental data with computational models but with limited accuracy. We developed a novel concept of comprehensive influence of proteins in network for reconstructing the PPI network. Our model outperforms the state-of-the-art ones and we then applied our model to identify novel interactions for further validation.

## 1. Introduction

Proteins are biological functional units, and protein-protein interaction (PPI) networks are associated with biological signal transduction, gene-expression regulation, energy and substance metabolism, and cell cycle regulation (Keskin, et al., 2016). Network reconstruction and analysis of disease associated PPIs is important to the understanding of protein functions in networks and of the molecular mechanism of diseases (Alvarez, et al., 2016; Azevedo, et al., 2018). The rapid development of experimental techniques to detect protein interactions has greatly increased the data accumulation for the reconstruction and analysis of PPI network; however, due to the limitations of experimental methods and the complexity of biomedical systems, the experimental results often present high false-positive and false-negative rates (Cheng, et al., 2016). Moreover, experimental verification of the PPI network is time consuming and expensive. Therefore, rapid and accurate computational methods represent useful alternative for predicting protein interactions, with such results providing guidance for experiments, including determination of unknown relationships between disease-causing genes and protein interactions (Zhu, et al., 2015). Such knowledge can be useful to the understanding of protein structure and evolution, as well as the overall function of the network and its associated dynamic processes (Cowen, et al., 2017; Hoeng, et al., 2014).

The unknown interaction between proteins can be predicted by the theory of complex network. PPI network consists of nodes and edges, with the nodes representing proteins and the edges representing associations or interactions between the two proteins (Gani, et al., 2015). Libennowell et al. considered the influence of common neighbor nodes in the network and introduced similarity indices to include common neighbors (Libennowell and Kleinberg, 2007). Pujari et al. suggested that each attribute of a pair of connected proteins represents different information (Pujari and Kanawati, 2012), and Li et al. proposed the use of a feature set containing user proximity, attributes, and topology for predictions (Li, et al., 2014). When analyzing protein interaction pairs, the above mentioned methods often take the local or path information of the nodes in the network into account, and do not systematically analyze the edge relationships in the network.

In the present study, we aim to emphasize the analysis of edges in the PPI network to improve the reconstruction of disease associated PPI network and develop the concept of network influence to evaluate the possibility of interactions between proteins. The model built based on the concept was then compared with the existed methods and was applied in reconstructing several disease associated PPI networks.

## 2. Results

### 2.1 Baseline and Evaluation Metrics

To evaluate the performance of our proposed method, six existed methods are used for comparison, they are structural deep-network embedding (SDNE) index (Crichton, et al., 2018; Wang, et al., 2016), Common Neighbors (CN) index (Libennowell and Kleinberg, 2007), Jaccard index (Gunes, et al., 2016), Sorenson index (Martinez, et al., 2017), HPI index (Martinez, et al., 2017) and Salton index(Martinez, et al., 2017).

To measure the performance of an algorithm, it is necessary to divide the data set into training set and test set. In link prediction, we remove 15% of the edges (retaining the related nodes) as the test set. The remaining 85% of the edges and all the nodes of the network are the training set. If no edges exist between two nodes, we define them as non-existent.

In this work, we used the AUC index to evaluate the performance of the methods (Lu, et al., 2009). AUC index measures the accuracy as a whole, and it refers to the probability that the score value of a random selected edge is higher than that of a non-existent edge in the test set. In the experiment, it get a similar score between each pair of nodes (edges in train set, test set and non-existent set) in the network by training set with an algorithm. One edge is randomly selected from the test set and another from the nonexistent edge. Among the *n* independent comparisons, if there are *n*′ cases that the edge in the test set has a higher score, then 1 point is added; if there are *n*″ cases that the edge in the test set and that of the nonexistent edge shares the same score, 0.5 point is added. Then the AUC index is defined as follows (Zhou, et al., 2009).

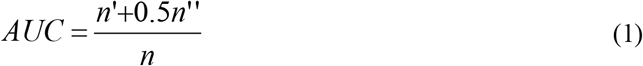

### 2.2 Results

Based on the perspective of graph theory, proteins and interactions in PPI network can be regarded as nodes and links respectively. The computational challenge about reconstructing PPI network from fixed seed proteins is referred as link prediction problem (Lei and Ruan, 2013). In this section, we present our results about the solution of the link prediction problem.

We first divided the data set into training set and test set (Hashemifar, et al., 2018). The basic format for each record of the data set involved in training and testing is <Protein A, Protein B, Interaction Score>. We take a certain proportion of the whole data set (e.g. 85%) as training set, and then set the record format of the rest (e.g. 15%) to < Protein A, Protein B, whether there is interaction >. After training, we input < Protein A, Protein B > in the test set to determine the interaction between protein A and protein B, and then compare the results with ground truth (Wang, et al., 2016).To evaluate the performance of the algorithms, we use our proposed method to construct interaction networks with five datasets as Table 1. In this section, we take 85% of each data set as training set and 15% as test set.

**Table 1.**
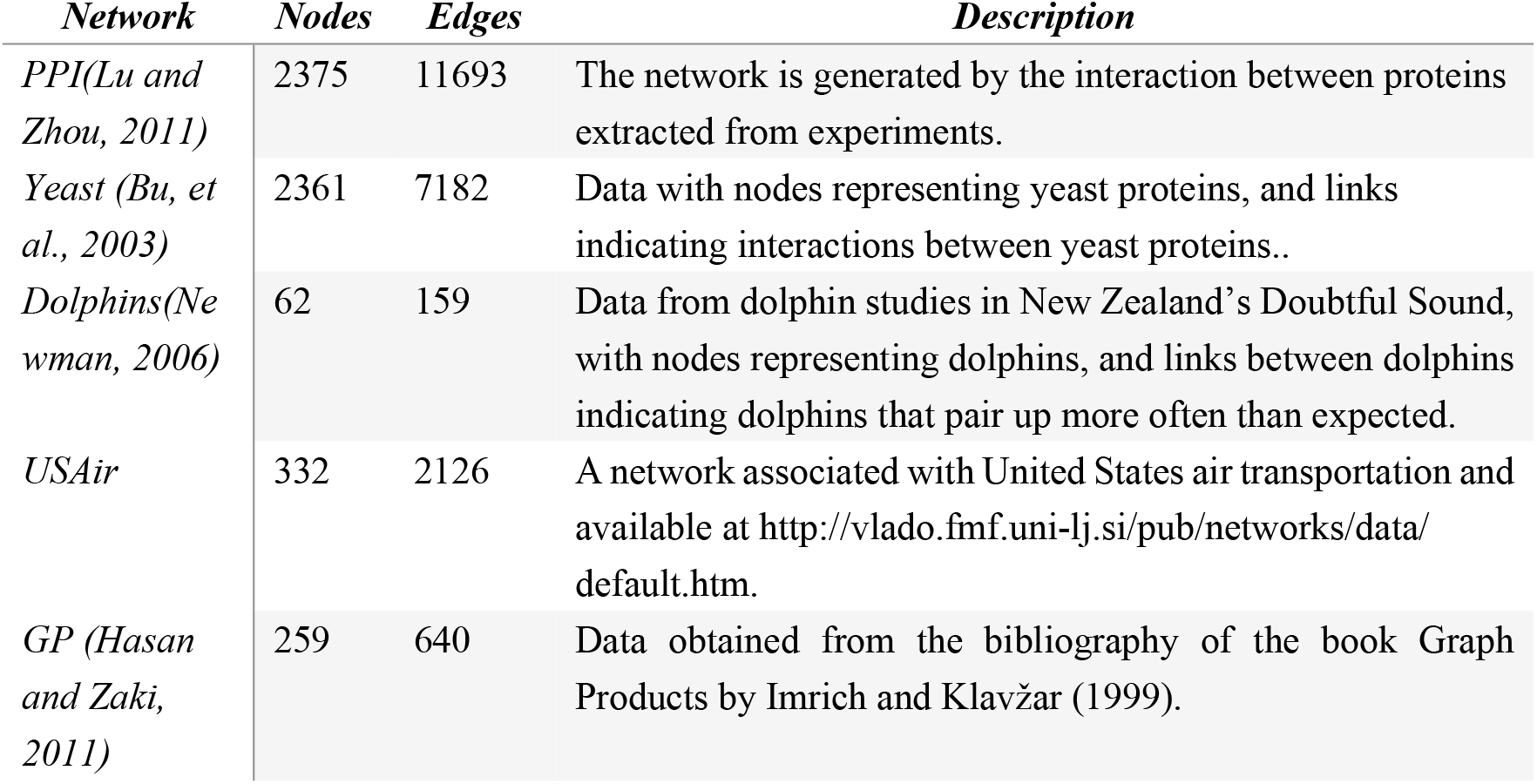
Interaction networks used for comparative analyses.

Table 2 shows that area under curve (AUC) of the proposed algorithm was improved relative to previous methods. We can get the following results from five different data sets: (1) In all data sets, the AUC value of our method is greater than 0.71, and the maximum value is more than 0.92, which proves that our method is effective in predicting the relationship between two nodes. (2) The AUC value of proposed method is improved compared with other six methods in five independent data sets. Only in the Dolphin network, the AUC value of our method is 0.003 lower than that of the CN method. The result proves that our method is superior to the other six methods. Fig 1 shows ROC curves of different networks with various methods. It can be seen that our method is effective for various network characterization and analysis.

**Fig 1.**
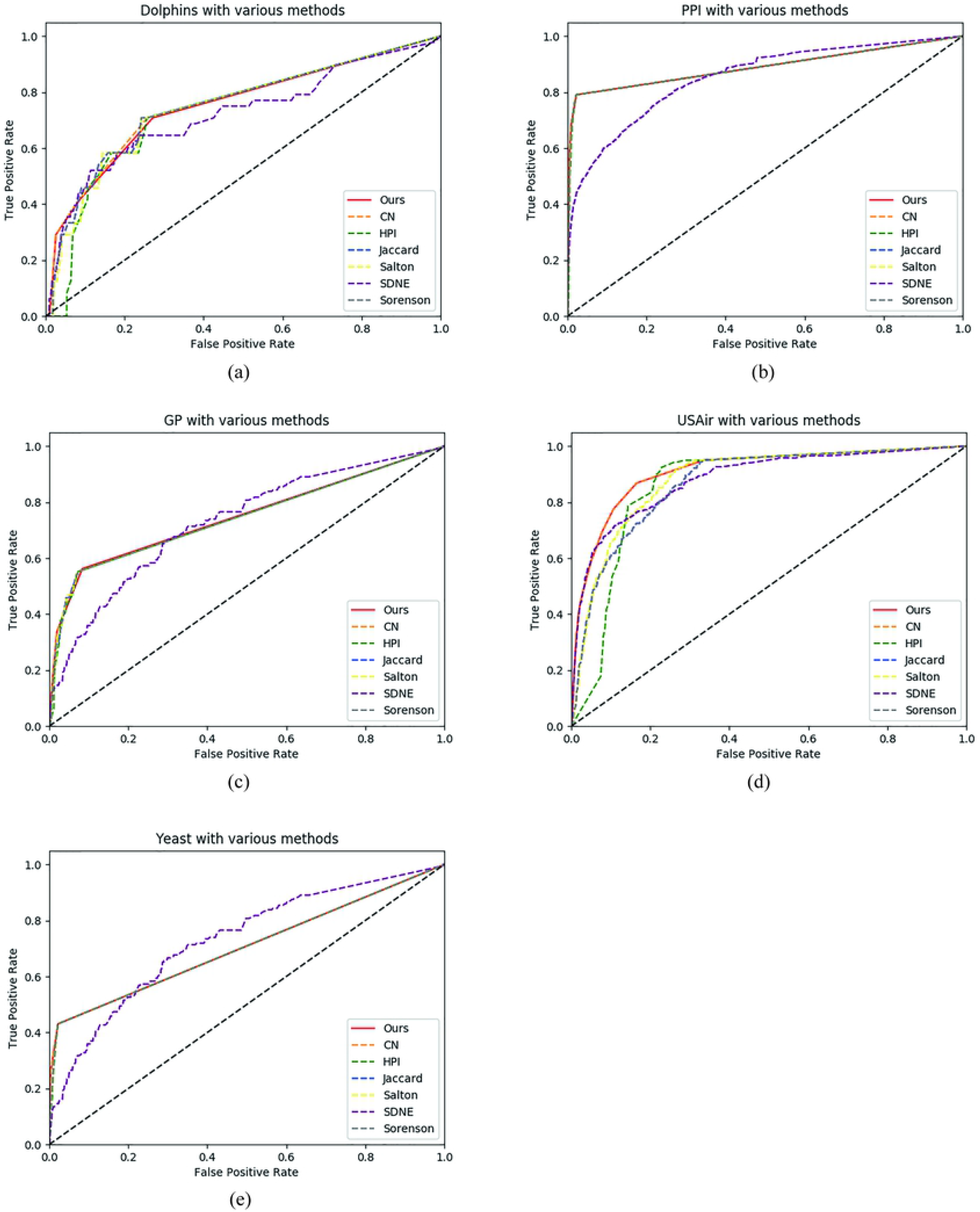
Performance of various methods for different network analysis.

**Table 2.** AUC of comparative analyses.

| *Dataset* | *Ours* | *SDNE* | *CN* | *Jaccard* | *Sorenson* | *HPI* | *Salton* |
| --- | --- | --- | --- | --- | --- | --- | --- |
| *PPI* | **0.8907** | 0.8868 | 0.8906 | 0.8892 | 0.8887 | 0.8071 | 0.8899 |
| *Yeast* | **0.7148** | 0.7090 | 0.7147 | 0.7076 | 0.7082 | 0.6002 | 0.7135 |
| *Dolphins* | 0.7748 | 0.7678 | **0.7778** | 0.7741 | 0.7740 | 0.7113 | 0.7700 |
| *USAir* | **0.9263** | 0.9163 | 0.9261 | 0.8865 | 0.8871 | 0.7881 | 0.8989 |
| *GP* | **0.7529** | 0.7355 | 0.7526 | 0.7451 | 0.7459 | 0.6394 | 0.7515 |

## 3. Discussion

In this section, we applied our method to reconstruct protein interaction networks and to identify potential interactions associated with prostate cancer and Parkinson’s disease.

### 3.1 Data Sources

We used the Human Protein Reference Database (HPRD) (Peri, et al., 2003) and IntAct Molecular Interaction Database as the data sources, the STRING database of known and predicted PPIs was used to verify our results (Jensen, et al., 2009).

The HPRD integrates information pertaining to domain architecture, posttranslational modifications, interaction networks, and disease associations for each protein in the human proteome. The proteins and interactions in the HPRD are manually extracted from the literature by experts after interpretation and analysis using domain knowledge. The IntAct Molecular Interaction Database houses PPI data manually extracted from published literature and includes experimental methods, conditions, and interacting domains, as well as information concerning non-interacting proteins.

We extracted data from the HPRD and IntAct to gather seed proteins related to prostate cancer and Parkinson’s disease, and then unified the various identifiers from multiple databases. After that, we collected the PPI dataset using the seed-protein data with the nearest-neighbor method and initialized the score of each protein-interaction pair.

### 3.2 PPI network of prostate cancer

Prostate cancer is a malignancy in the urinary system that primarily targets middle-aged and elderly men, with an estimated 1,600,000 cases diagnosed and 366,000 deaths annually (Torre, et al., 2015). To identify genes and proteins related to cancer progression is important to the personalized treatment of this disease (Zhang, et al., 2016; Zhao, et al., 2017). To screen prostate cancer associated proteins, we set *γ* as 0.5 to calculate influence between two nodes, and set *β* as 0.5 to extract top influence.

#### 3.2.1 Seed-protein identification and unification

Using “prostate” as a retrieval term in the HPRD, we identified 18 proteins for the seed-protein set. We then used UNIPROT to identify prostate-cancer-related genes according to standard gene identifiers, which were subsequently mapped to the seed-protein set according to Swiss-Prot protein identifiers (Table 3).

**Table 3.** Prostate cancer seed-protein set.

|  | ***HPRDID*** | ***SWISSPROTID*** | ***PROTEIN*** |
| --- | --- | --- | --- |
| **1** | 05084 | O96017 | CHK2 checkpoint homolog |
| **2** | 01143 | P08118 | Beta-microseminoprotein |
| **3** | 02437 | P10275 | Androgen receptor |
| **4** | 01885 | P12830 | Cadherin 1 |
| **5** | 01092 | P21757 | Macrophage scavenger receptor 1 |
| **6** | 00625 | P22455 | Fibroblast growth factor receptor 4 |
| **7** | 02997 | P29323 | EphB2 |
| **8** | 08926 | P35680 | Hepatocyte nuclear factor 1-beta |
| **9** | 02486 | P50539 | Max interacting protein 1 |
| **10** | 02554 | P51587 | BRCA2 |
| **11** | 03431 | P60484 | Protein tyrosine phosphatase PTEN |
| **12** | 01590 | Q05823 | Ribonuclease L |
| **13** | 00075 | Q15911 | Alpha fetoprotein enhancer binding protein |
| **14** | 10109 | Q8NDI1 | EH domain-binding protein 1 |
| **15** | 05210 | Q92826 | Homeobox protein Hox-B13 |
| **16** | 03632 | Q99612 | Core promoter element binding protein |
| **17** | 05641 | Q9BQ52 | ELAC |
| **18** | 04065 | Q9Y6D9 | MAD1 mitotic arrest deficient-like 1 |

#### 3.2.2 Establishment of prostate related PPI network

We collected interacting-protein pairs related to prostate cancer according to our generated seed-protein set. To assess the confidence of each interaction, we used the following scoring rules: 1) protein pairs were assigned a confidence value of 0.8 from the HPRD and 2) other protein pairs were assigned a confidence value from IntAct. This allowed identification of the PPI dataset from the seed-protein set via the nearest-neighbor method, resulting in 488 protein pairs.

The constructed PPI network contained 450 nodes, with an average degree of 2.19 for each node. The network comprised a weakly connected graph that met the characteristics of a small-world network (Fig 2).

**Fig 2.**
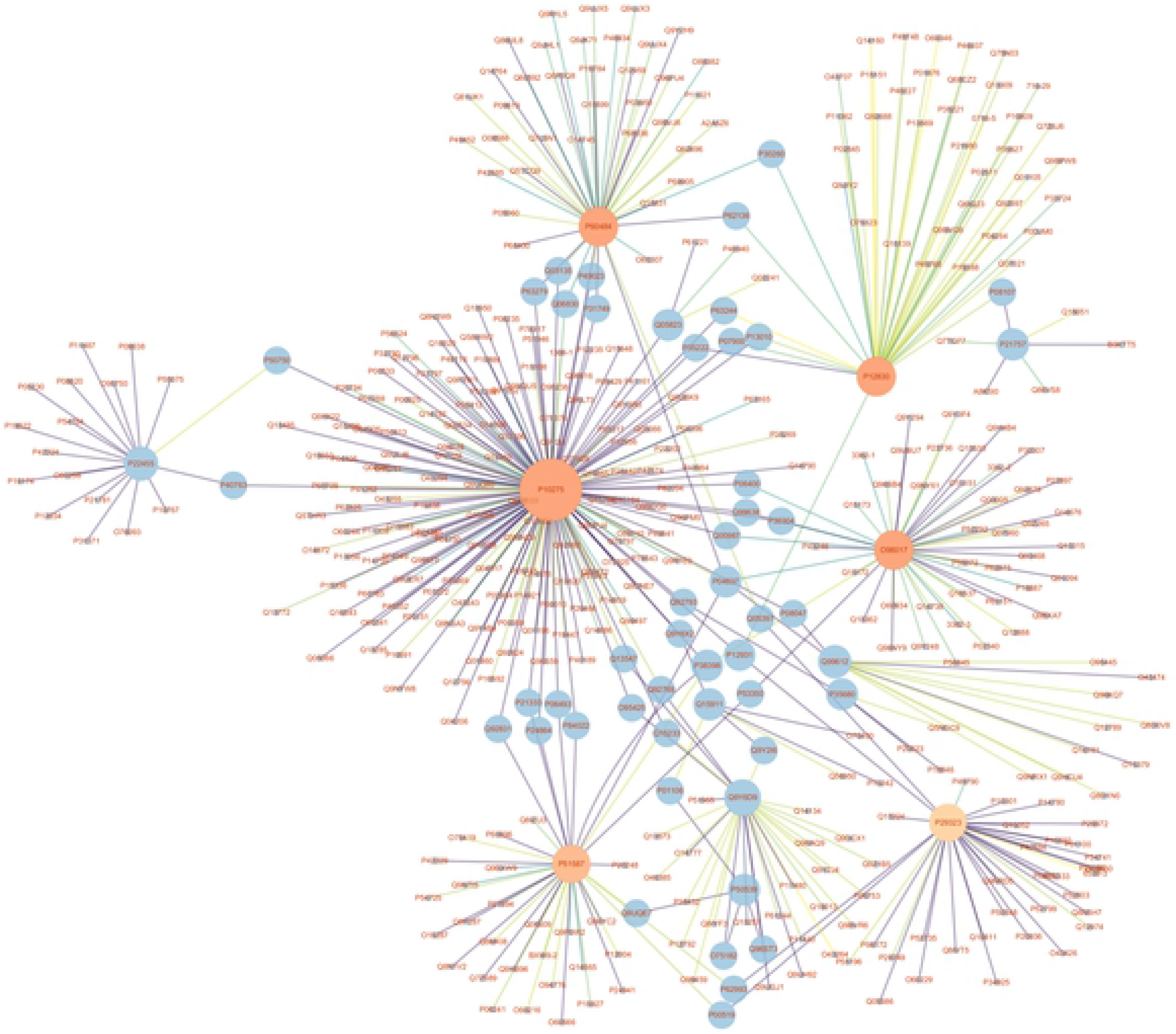
Overview of the prostate cancer PPI network.

Generation of the scale-free graph resulted in hub nodes associated with the proteins CDH1, EPHB2, BRCA2, PTEN, and CHEK2, with their importance in maintaining network stability suggesting their potentially key roles in the pathogenesis of prostate cancer. Upregulated proteins in tumors likely play a role in tumor invasion and metastasis, with examples including TP53, MDM2, RB1, HDAC1, HDAC2, and SRC. We use cytoscape (Shannon, et al., 2003) to visualize the PPI network, and Fig 2 shows the PPI network associated with prostate cancer, and Fig 3 shows details of specific high-degree hubs, including those associated with AR, CDH1, EPHB2, BRCA2, PTEN, and CHEK2.

**Fig 3.**
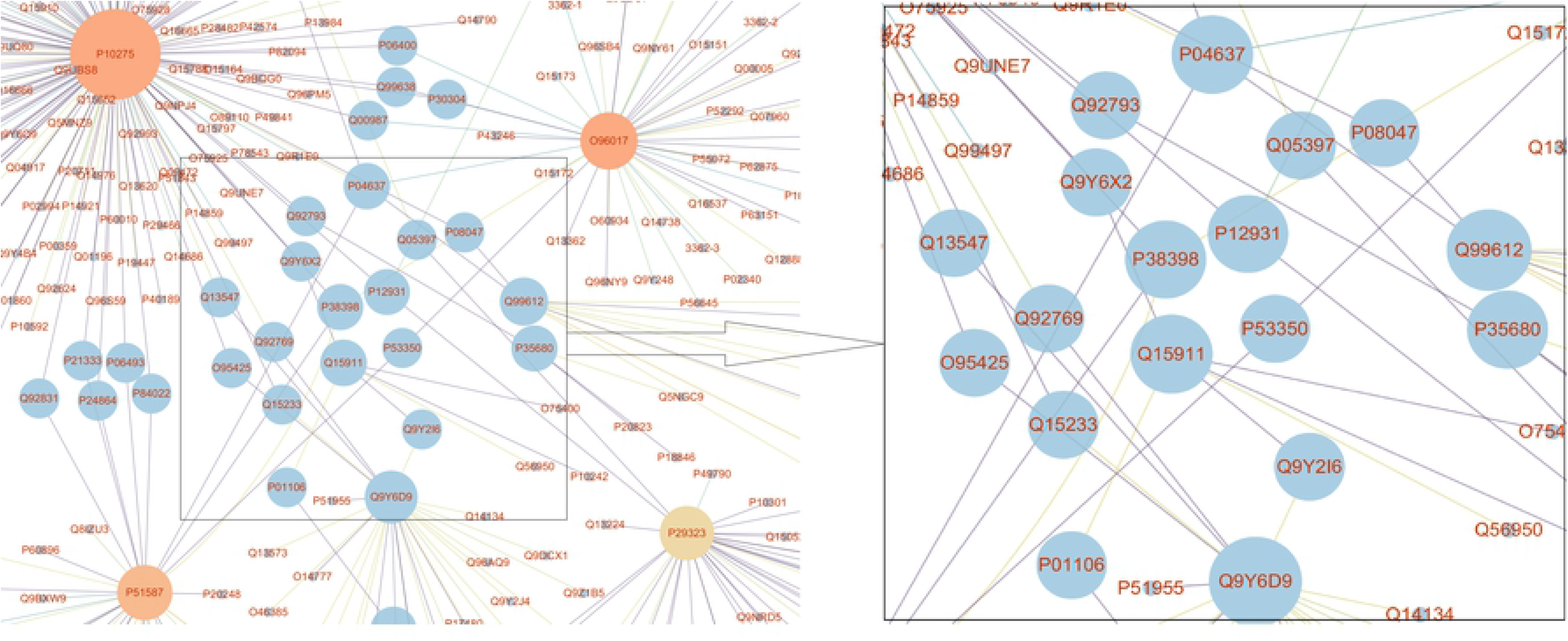
Representative central nodes. Nodes included AR(P10275), CDH1(P12830), and CHEK2(O96017) connected with several hub nodes in the network.

We choose top 24 proteins from established PPI network in Fig 3 (as listed in Table 4). These proteins are highly connected proteins in the PPI network associated with prostate cancer, which are closely connected with several hub nodes of the network, and it is important to maintain the stability of the network.

**Table 4.** Highly connected proteins in the PPI network associated with prostate cancer.

|  | *GENE* | *SWISSPROTID* | *PROTEIN* |
| --- | --- | --- | --- |
| **1** | KAT2B | Q92831 | PCAF |
| **2** | FLNA | P21333 | Filaminranks A, alpha |
| **3** | CDK1 | P06493 | CDC2 |
| **4** | CCNE1 | P24864 | Cyclin E1 |
| **5** | SMAD3 | P84022 | SMAD family member 3 |
| **6** | GRB2 | P62993 | Grb2 |
| **7** | ABL1 | P00519 | v-abl Abelson murine leukemia viral oncogene homolog 1 |
| **8** | SVIL | O95425 | Supervillin |
| **9** | HDAC1 | Q13547 | Histone deacetylase 1 |
| **10** | HDAC2 | Q92769 | Histone deacetylase 2 |
| **11** | NONO | Q15233 | Non pou domain containing octamer binding protein |
| **12** | SRC | P12931 | c-Src |
| **13** | CDC27 | P30260 | Anaphase promoting complex, subunit 3 |
| **14** | PPP1CA | P62136 | Protein phosphatase 1, catalytic subunit, alpha isoform |
| **15** | CAV1 | Q03135 | Caveolin 1 |
| **16** | PXN | P49023 | Paxillin |
| **17** | UBE2I | P63279 | Ubiquitin conjugating enzyme E2I |
| **18** | PRDX1 | Q06830 | Peroxiredoxin 1 |
| **19** | AKT1 | P31749 | AKT1 |
| **20** | RB1 | P06400 | Retinoblastoma 1 |
| **21** | RAD9A | Q99638 | RAD9 |
| **22** | CDC25<br>A | P30304 | CDC 25A |
| **23** | MDM2 | Q00987 | MDM2 |
| **24** | TP53 | P04637 | p53 |

#### 3.2.3 Prostate cancer associated network analysis

The analysis of prostate cancer associated PPI network indicates that only a small portion of the proteins in the network were highly connected, with most having short-path connections which is in agreement with the characterization of a small-world network. Additionally, these results are consistent with the characteristics of PPI networks associated with complex diseases, in that the network extends from a central node to the periphery. In most cases, the node degree of a complex network obeys a power-law distribution rather than a Poisson distribution (Virkar, 2014). For a randomly selected node, the probability of its degree being *k* is *P*(*K*) ∝ *K^r^*, where *r* and *k* are constants, thereby representing a scale-free network. Previous studies suggest that many complex networks are characterized by scale-free properties. Similarly, our analysis indicated that the generated network displayed a node-degree distribution representative of a scale-free network (Fig 4).

**Fig 4.**
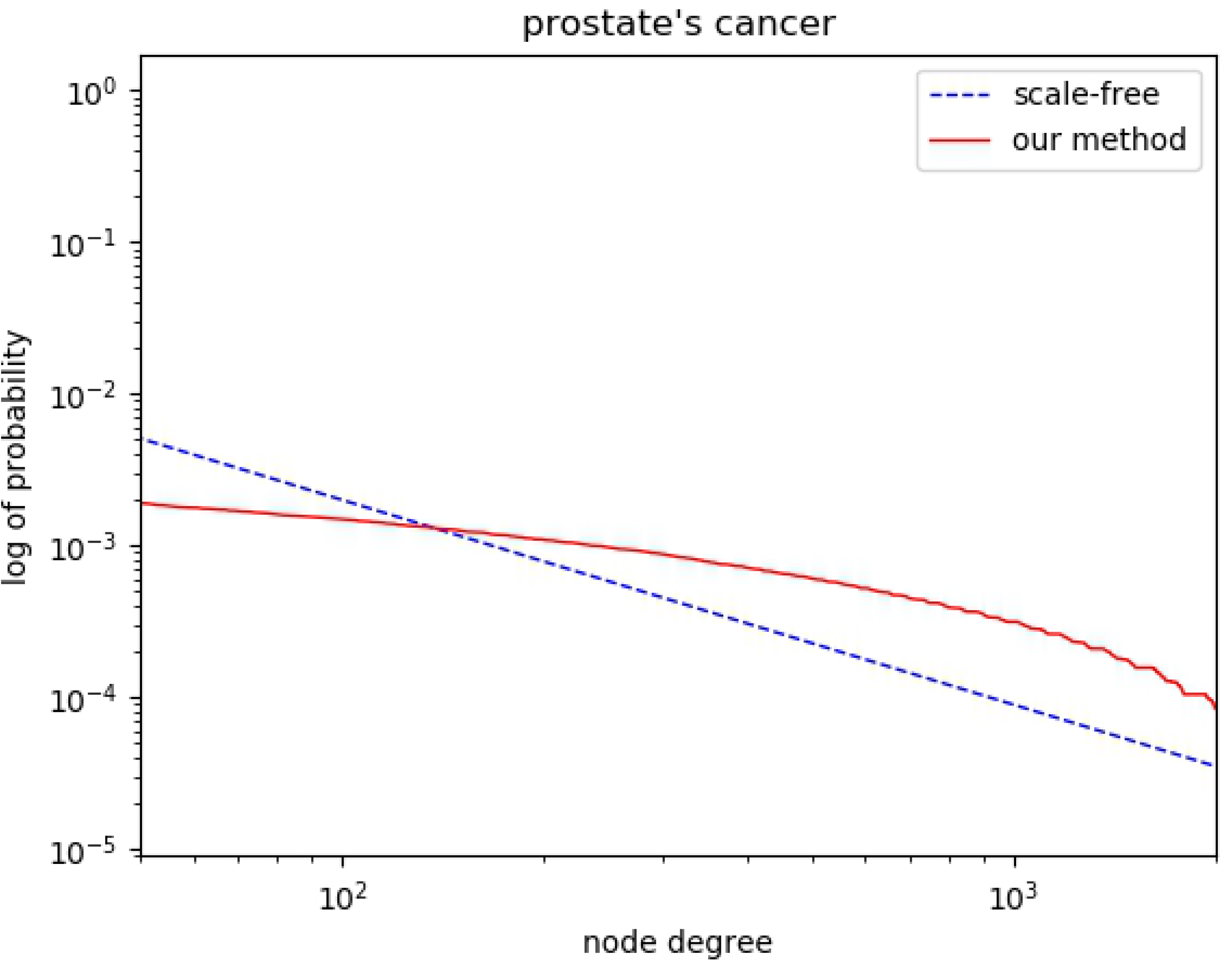
Log of the probability of the node-degree distribution of the PPI network associated with prostate cancer and representative of a scale-free network.

We selected proteins with the highest influence scores, resulting in an average of 75% confirmed PPIs with those in STRING, the other 25% verified in the literature. We identified two proteins with novel interactions (AQP1 and MMP2). Studies suggest that AQP1 upregulates MMP2 and MMP9 levels, leading to increased invasion and adhesion of colon cancer cells and migration of bladder cancer cells. Aquaporins (AQP) are widely distributed in tissues and cells, with AQP1 an integrative membrane protein in human erythrocytes responsible for maintaining cell permeability. Urine proteome analysis to identify disease biomarkers for prostate cancer, renal cell carcinoma, bladder cancer, urothelial carcinoma, and renal Fanconi syndrome suggested as a possible marker AQP1 (Morrissey, et al., 2015; Morrissey, et al., 2014; Rodrigues, et al., 2017; Rubenwolf, et al., 2009; Tomita, et al., 2017).

MMP2 is a protease capable of degrading collagen and the extracellular matrix and involved in tumor-cell infiltration of connective tissue stroma, small blood vessels, and lymphatic tissue (Wu, et al., 2019). Additionally, MMP2 is highly expressed in prostate cancer tissues and plays a role similar to that of proto-oncogenes, as attenuated MMP2 levels inhibit tumor-cell proliferation, invasion, and migration and promote apoptosis, thereby alleviating tumor malignancy.

Relationships between AQP1 and prostate cancer invasion and metastasis in connection with MMP2 status remain unknown; however, previous studies implicated both AQP1 and MMP2 are associated with prostate cancer (Brundl, et al., 2018; Lacorte, et al., 2015). AQP1 is localized to the kidney, with studies reporting its elevated expression in the prostate gland in testicles of rats. Additionally, Kyoto Encyclopedia of Genes and Genomes (KEGG) analysis indicated that MMP2 is involved in bladder cancer and other cancer pathways. Although investigations have focused on their roles in colon cancer (Knight, et al., 2016; Yde, et al., 2016) and *in vivo* status in animal studies (Jiang, et al., 2019), the relationship between these two proteins and prostate cancer is not reported yet. Our analysis suggested a potential role for these two proteins in prostate cancer; although experimental studies are necessary to confirm this relationship.

### 3.3 PPI network of Parkinson’s disease

Parkinson’s disease (Berg, et al., 2018) is a common age-related brain disorders defined primarily as a movement disorder accompanied by typical symptoms of resting tremors, rigidity, bradykinesia, and postural instability (Aarsland, et al., 2017). For Parkinson’s disease, we detected indirect influence have negative effects on results with the same parameter as that for prostate cancer. So *γ* was set as 0.45 to reduce the indirect impact, and *β* was set as 0.05 to improve accuracy in top.

#### 3.3.1 Seed-protein identification and unification

Using Parkinson’s disease as retrieval word in the HPRD, we obtained 14 proteins for the seed-protein set and mapped them to gene symbols and Swiss-Prot protein identifiers (Table 5).

**Table 5.** Parkinson’s disease seed-protein set.

|  | ***GENE*** | ***HPRID*** | ***SWISSPROTID*** | ***PROTEIN*** |
| --- | --- | --- | --- | --- |
| **1** | MAPT | 01142 | P10636 | Microtubule associated protein tau |
| **2** | SNCA | 01227 | P37840 | Synuclein alpha |
| **3** | TRPM7 | 10418 | Q96QT4 | Transient receptor potential cation channel, subfamily M, member 7 |
| **4** | PINK1 | 10514 | Q9BXM7 | PTEN induced putative kinase 1 |
| **5** | HTRA2 | 05919 | O43464 | HtrA serine peptidase 2 |
| **6** | PARK7 | 03961 | Q99497 | Oncogene DJ1 |
| **7** | PARK2 | 03967 | O60260 | Parkinson disease (autosomal recessive, juvenile) 2, parkin |
| **8** | PRKAG2 | 04119 | Q9UGJ0 | PRKAG2 |
| **9** | SNCAIP | 04804 | Q9Y6H5 | Synphilin 1 |
| **10** | TAF1 | 02436 | P21675 | Transcription factor IID |
| **11** | NDUFV2 | 02757 | P19404 | NADH ubiquinone oxidoreductase flavoprotein 2 |
| **12** | NR4A2 | 03493 | P43354 | Nuclear receptor subfamily 4, group A, member 2 |
| **13** | UCHL1 | 01877 | P09936 | Ubiquitin carboxyl terminal esterase L1 |
| **14** | ADH1C | 00066 | P00326 | Alcohol dehydrogenase 3 |

#### 3.3.2 Establishment of Parkinson’s disease PPI network

Application of the scoring rules described above resulted in identification of 568 pairs of interacting proteins associated with Parkinson’s disease using the nearest-neighbor method.

The average node degree in the PPI network was 2.12, which fit the characteristics of a small-world network (Fig 5). Fig 5 shows the PPI network associated with Parkinson’s disease, and Fig 6 shows details of representative central nodes including SNCA, PARK2, MAPT connected with several hub nodes in the network.

**Fig 5.**
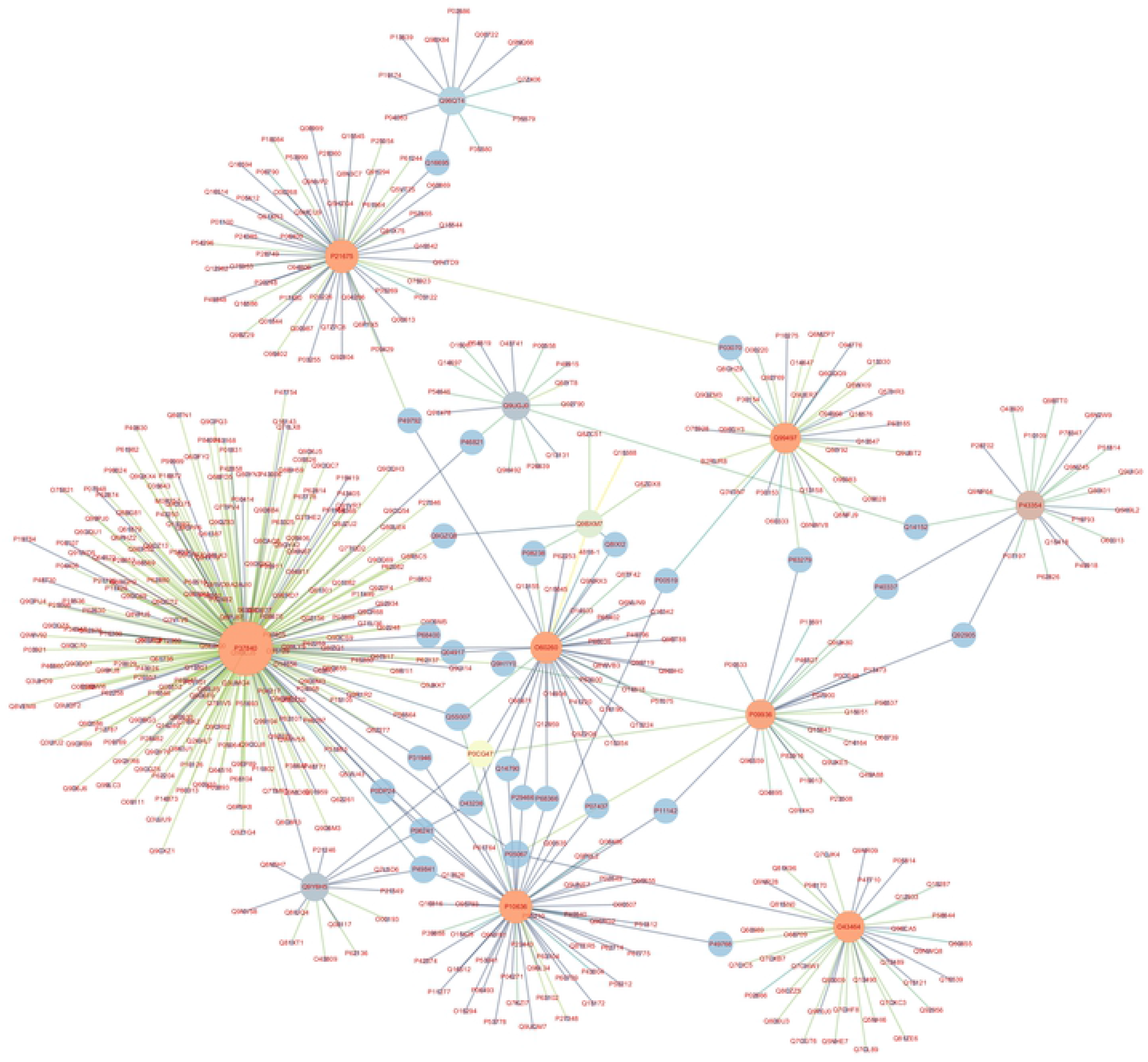
Overview of the Parkinson’s disease PPI network.

We choose top 21 highly connected proteins from the PPI network associated with Parkinson’s disease. We can see that these proteins are almost identical to the blue proteins in Fig 6. These proteins maintain the stability of the network and are connected with several hub nodes of the network. Proteins such as UBB, HSPA8, ABL1 and CASP8 are involved in biological processes of Parkinson’s disease, for example, UBB shows relationship with positive regulation of protein monoubiquitination (Kazlauskaite, et al., 2014) and HSPA8 is related to regulation of protein import and regulation of protein complex assembly (Orenstein and Cuervo, 2010). LRRK2 may play a role in the phosphorylation of proteins central to Parkinson’s disease, which has certain effects on biological processes and molecular functions (Angeles, et al., 2011). Table 6. Shows highly connected proteins in the PPI network associated with Parkinson’s disease.

**Fig 6.**
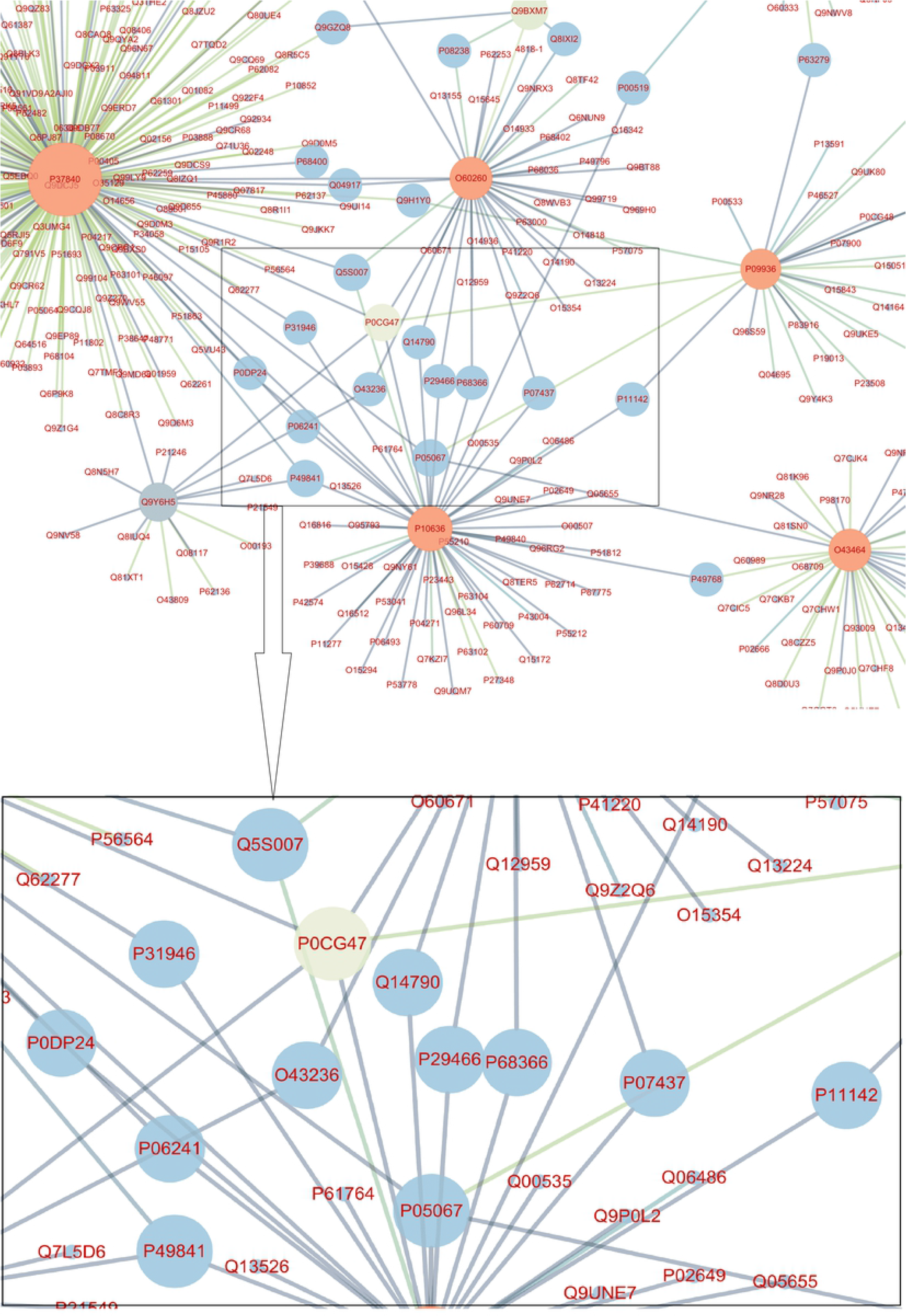
Representative central nodes in the Parkinson’s disease PPI network. Nodes including SNCA(P37840), PARK2(O60260), MAPT(P10636) connected with several hub nodes in the network.

**Fig 7.**
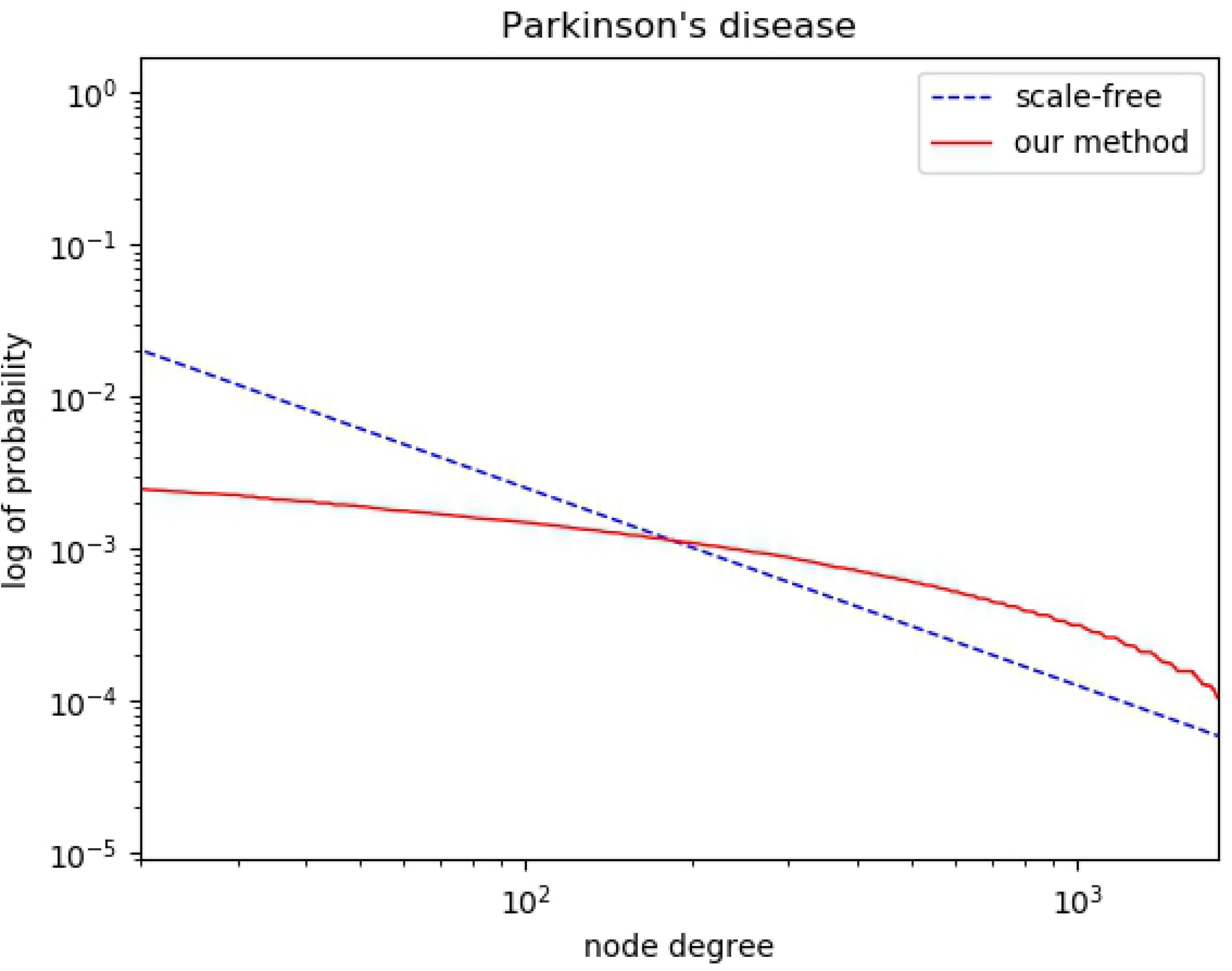
Log of the probability of the node-degree distribution of the PPI network associated with Parkinson’s disease and representative of a scale-free network.

**Table 6.** Highly connected proteins in the PPI network associated with Parkinson’s disease.

|  | *GENE* | *HPRID* | *SWISSPROTID* | *PROTEIN* |
| --- | --- | --- | --- | --- |
| **1** | UBB | 06771 | P0CG47 | Ubiquitin B |
| **2** | LRRK2 | 10613 | Q5S007 | Leucine rich repeat kinase 2 |
| **3** | APP | 00100 | P05067 | Amyloid beta A4 protein |
| **4** | GSK3B | 05418 | P49841 | Glycogen synthase kinase 3 beta |
| **5** | FYN | 00655 | P06241 | Fyn |
| **6** | YWHAB | 03184 | P31946 | 14-3-3 Beta |
| **7** | SEPT4 | 04738 | O43236 | Septin 4 |
| **8** | CASP8 | 03459 | Q14790 | Caspase 8 |
| **9** | CASP1 | 00977 | P29466 | Caspase 1 |
| **10** | TUBA4A | 01851 | P68366 | Tubulin alpha 1 |
| **11** | TUBB | 01852 | P07437 | Tubulin, beta |
| **12** | HSPA8 | 07205 | P11142 | Heat shock 70 kDa protein 8 |
| **13** | PSEN1 | 00087 | P49768 | Presenilin 1 |
| **14** | YWHAH | 00215 | Q04917 | 14-3-3 Eta |
| **15** | ATG5 | 16051 | Q9H1Y0 | Autophagy protein 5-like |
| **16** | MAP1LC3B | 14358 | Q9GZQ8 | Microtubule-associated protein 1 light chain 3 beta |
| **17** | HSP90AB1 | 06763 | P08238 | HSP90B |
| **18** | RHOT1 | 15247 | Q8IXI2 | Mitochondrial Rho 1 |
| **19** | ABL1 | 01809 | P00519 | v-abl Abelson murine leukemia viral oncogene homolog 1 |
| **20** | UBE2I | 09045 | P63279 | Ubiquitin conjugating enzyme E2I |
| **21** | RANBP2 | 03111 | P49792 | Ran binding protein 2 |

#### 3.3.3 Parkinson’s disease associated network Analysis

As shown in Fig. 8, the node-degree distribution in the Parkinson’s disease associated network also met the characteristics of a scale-free network, which conforms to a power-law distribution. For the Parkinson’s disease data, 69% of the predicted PPI can be validated by STRING. The others are the novel Parkinson’s disease associated PPI and are listed in Table 7.

**Fig 8.**
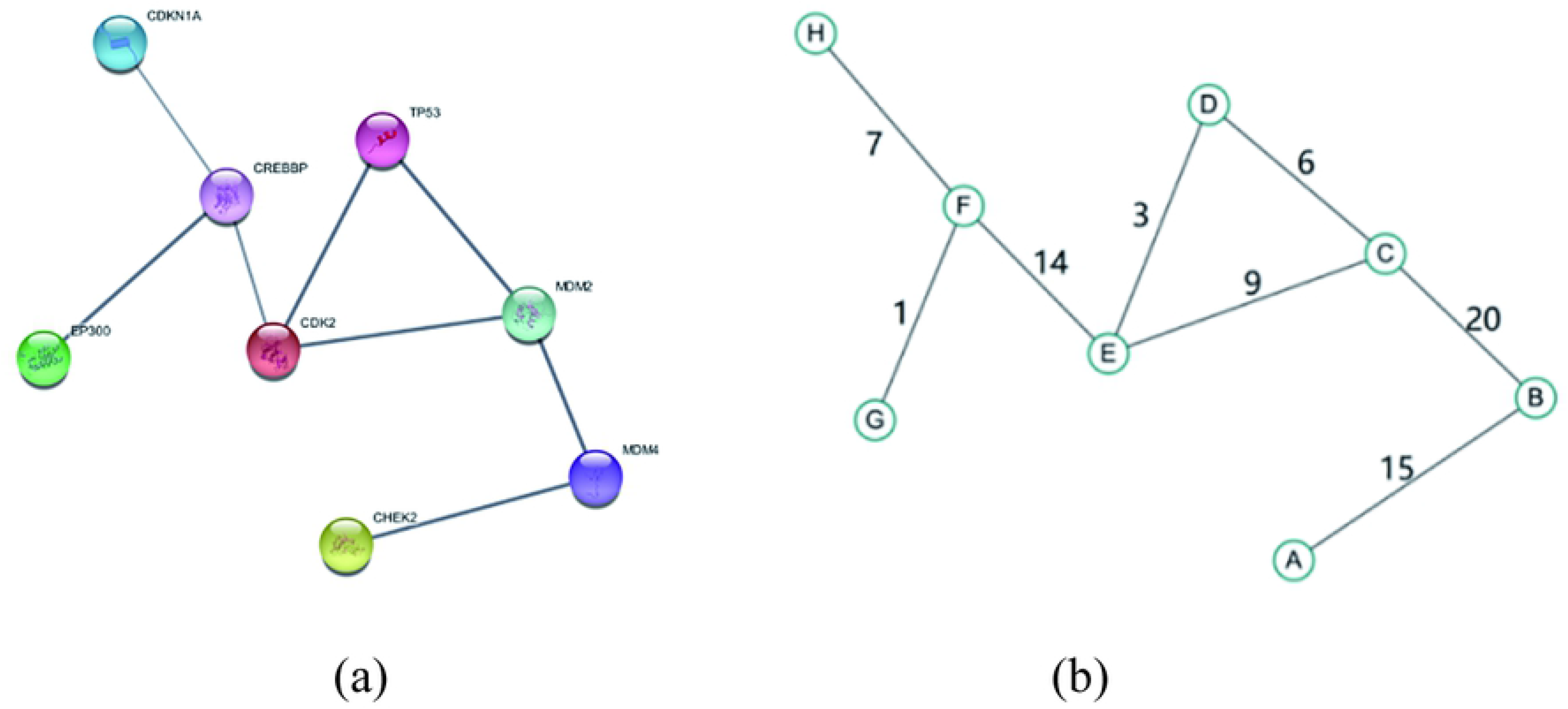
(a) is part of the TP53 associated PPI network and (b) is the weighted PPI network abstracted from the left.

**Table 7.**
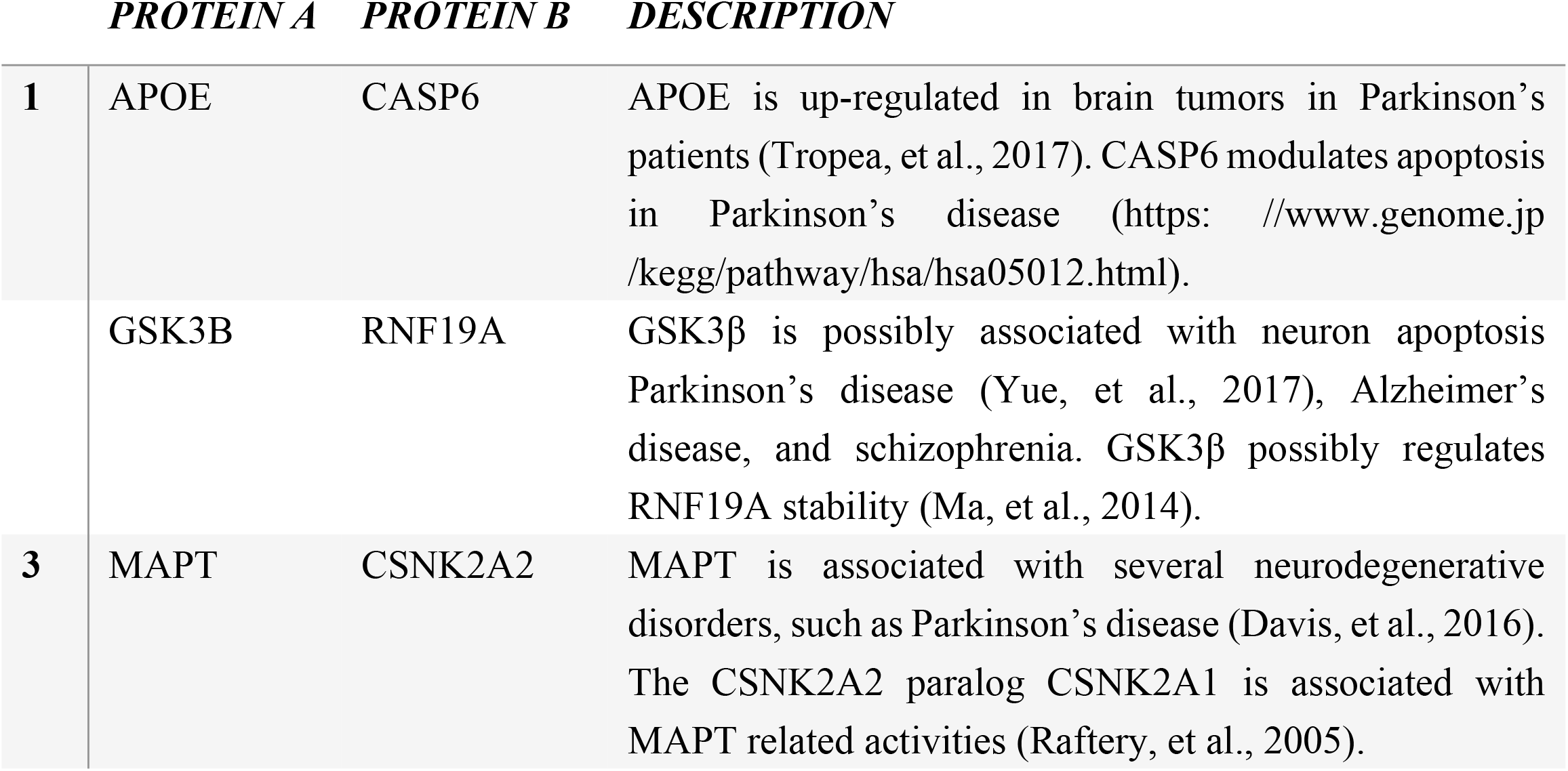
Novel PPIs identified using our proposed method.

We investigated the functions of the identified interacting proteins with the highest influence scores (APOE, CASP6, GSK3B, RNF19A, MAPT, and CSNK2A2). The PPI network included several central nodes (TAF1, PARK7, SNCA, PARK2, UCHL1, MAPT, HTPR2, and NR4A2) involved in Parkinson’s disease (Campelo and Silva, 2017; Domingo, et al., 2016; Rocha, et al., 2018), as well as others (UBB, LRRK2, APP, GSK3B, FYN, SEPT4, RANBP2, ABL1, and MAP1LC3B).

Understanding complex biological processes involves high levels of data beyond the scope of experimental methods; therefore, computational approaches for reconstruction of disease associated PPI network have become increasingly important. Here, we compare our proposed methods with six existed methods and the results indicate that our method is effective to identify disease associated proteins and their interactions. We then generated two novel PPI networks associated with prostate cancer and Parkinson’s disease using our method. A novel interaction between AQP1 and MMP2 in prostate cancer and three novel potential interaction pairs for APOE and CASP6, GSK3B and RNF19A, MAPT and CSNK2A2 in Parkinson’s disease were predicted for further experimental validation. Our results suggest that this method could be applied to different diseases for identifying disease associated PPIs.

## 4. Materials and Methods

A PPI network can be described as an undirected graph, *G* = (*V, E*), where *V* denotes the set of nodes, and *E* represents the set of edges. The importance of the interaction or node in a network can be described by different wrights. Fig 8 shows an example of PPI network, where the left is a part of PPI network extracted from STRING database with TP53 as the keyword and then visualized in Cytoscape (Shannon, et al., 2003) and the right is the weighted PPI network abstracted from the left.

In a concrete PPI network, the function of a protein is influenced by both its direct and indirect interactions, therefore it is necessary to consider the effect of its surrounding sub-network. To evaluate the influence of a protein in a network, we introduce the following definitions, taking the network shown in Fig 8 as an example.

### Definition 1

<A, B> represents a direct influence and is used to evaluate direct interactions between two proteins, *A* and *B*.

For example, in Fig 8, the influence of <*A, B*> for nodes *A* and *B* is 15 (direct influence), and that of <*B*, *C*> is 20.

### Definition 2

Indirect influence is used to evaluate an indirect interaction between two proteins through other proteins.

For example, in Fig 8, nodes *A* and *B* exhibit only a direct influence (i.e., <*A*, *B*>), whereas we define node *A* as having an interaction with node *C* through node *B* (i.e., an indirect influence; <*A*, *B*, *C*>). Because indirect influence is exerted by other proteins, it is relevant to more than two primary nodes. In Fig 8, nodes *A* and *C* might indirectly influence one another through node *B*.

### Definition 3

An indirect influence involves the shortest distance between two nodes.

For example, in Fig 8, two pathways from *D* to F include <*D, E, F*> and <*D, C, E, F*>, respectively. When there are additional pathways between two proteins, for the purpose of simplification, we only consider the case with the shortest distance.

### Definition 4

A hop between nodes *i* and *j* is the minimum number of nodes required to traverse from node *i* to node *j* in the network.

We use the following approach to generate a score for each protein by evaluating its value and influence to other proteins in the PPI network:

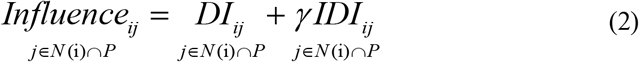

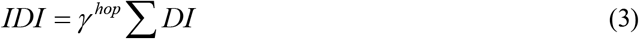

where *i* and *j* represent two solid nodes, *γ* is a discount(0<*γ*<1), *P* is the set of the protein network, and *N*(*i*) represents the set of proteins directly interacting with node *i*. In Equation (2), *Influence_ij_* denotes the influence between nodes *i* and *j* and is the sum of all direct and indirect influence scores. *DI_ij_* represents the set of direct links for nodes without considering the possible influence of shared nodes. In the case of indirect links, we assume that node *i* can possibly influence node *j* through node k, which is termed a “one-hop link”:

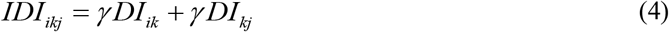

Because there are likely many one-hop links, there will consequently be many one-hop influence scores. In Equation (4), *k* is a variable node that links nodes *i* and *j*.

*IDI_iklj_* is used to evaluate the two-hop influence between nodes *i* and *j* and assumes that node *i* can get to node *j* through nodes *k* and *l*, and vice versa. For example, node *i* is linked to variable node *k*, which is linked to node *l*, and node *l* is linked to node *j*. We then assume that node *i* has possible influence on node *j* as part of a two-hop link:

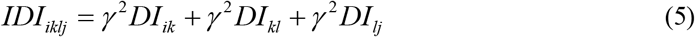

Link prediction based methods for predicting protein-protein interactions have their own merits, and data preprocessing has different effects on the performance of the algorithm. In this work, in order to reduce the problem caused by data bias, multiple data sources were used to optimize the parameters and maintain the data diversity for the generality of model. We proposed a link-based prediction method to extract protein interaction information from multiple data sources, and then evaluate candidate proteins. For the reconstruction of PPI network, proteins and links with high scores are selected based on Equation 1, as explained in Algorithm 1 and Algorithm 2.

Algorithm 1 describes the calculation of comprehensive influence between two nodes (i.e. proteins) in a network. Algorithms 2 describes how to build protein interaction networks based on the selected proteins from Algorithm 1.

**Algorithm 1:**
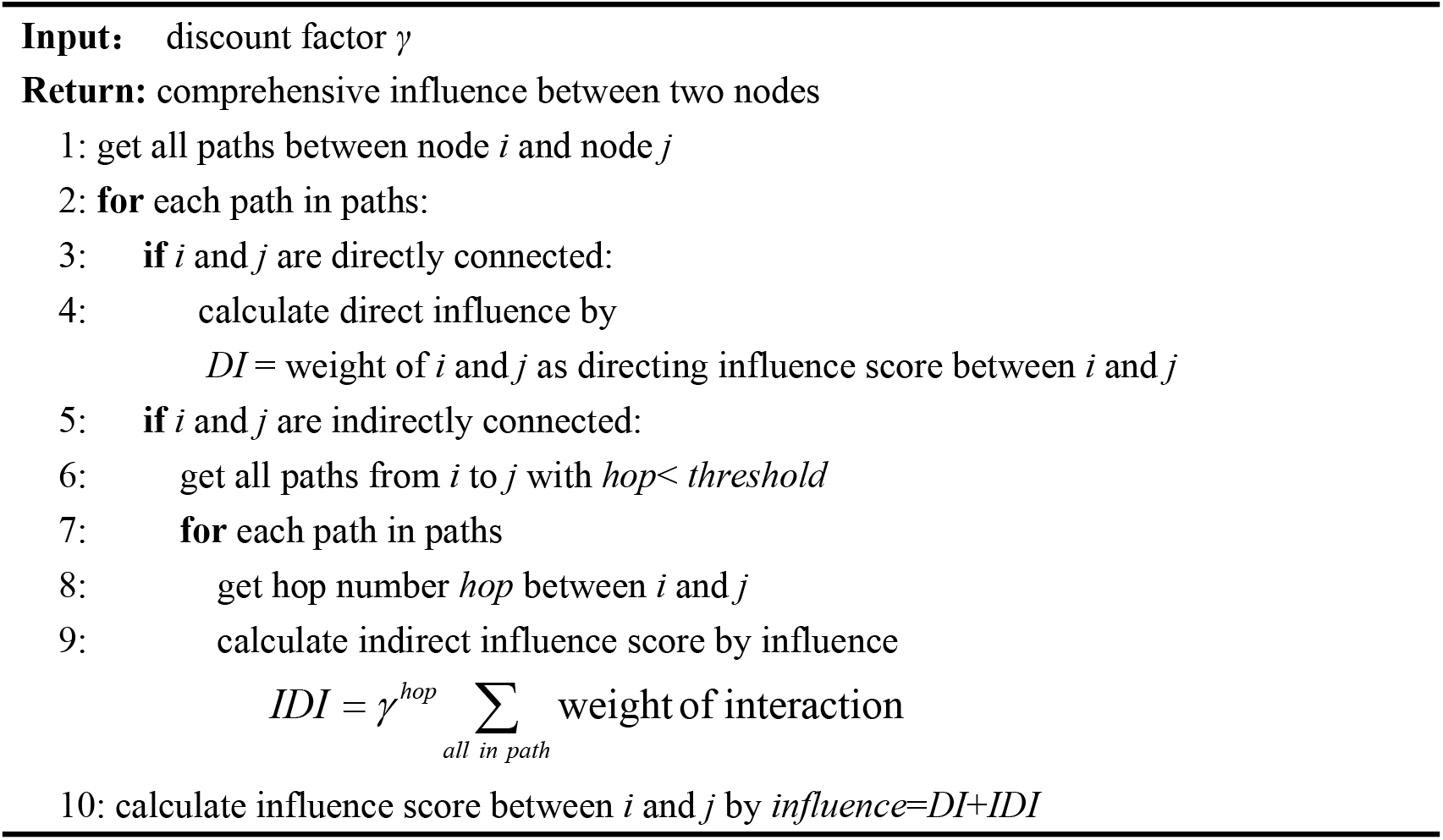
Computing comprehensive influence in the network

### Algorithm 2: Constructing PPI network via comprehensive influence

~~~
**Input:** interaction data, threshold *β*
**Return**: *G*
1: extract interactive data (node a, node b, weight)
2: initialize the score between interaction pairs
3: divide Train list and Test list
4: create the network G of relationships by retrieving direct partners from Train list
5: **for** each node in G
6:  calculate influence score between a and b with Algorithm 1
7:  sort candidate proteins of a in descending order
8:  acquire the top *β*% of the candidate proteins
9: **end for**
~~~

## 5. Related Works

Several databases containing protein-protein interactions or association data set are available such as, HPRD (Peri, et al., 2003), IntAct (Kerrien, et al., 2007), and STRING (Jensen, et al., 2009). However, previous studies suggested that high false-positive rate existed with much of the experimental data used to construct these networks, requiring improvements to yield more accurate PPI networks (Sze-To, et al., 2016). There are a number of online resources allowing access to interactome data (Szklarczyk and Jensen, 2015), including those for interactomes associated with *Arabidopsis* (Lee, et al., 2015) and maize (Zhu, et al., 2016), generated using predictive algorithms, experimental evidence, or a combination of the two. Although high-throughput experiments have allowed generation of large amounts of PPI data, issues concerning false-positive and false-negative results exist. Additionally, small-scale experiments displaying relatively high levels of accuracy require significant resources and manpower. Therefore, many computational methods have been proposed for protein interaction prediction to complement the experimental methods.

Computational reconstruction of PPI networks are based on different strategies, among which the similarity-based link prediction method is the most widely used. The basic idea is that if two nodes are similar, they are more likely linked. Based on the network resource allocation, the resource allocation index (Zhou, et al., 2009) and Adamix-Adar index (Adamic and Adar, 2003) are proposed and both of them take the degree information of network nodes into account, and regard the amount of resource transfer information between two nodes as similarity index. Based on the common neighbors index (Libennowell and Kleinberg, 2007), the number of common neighbors is used as a measure of similarity. Considering the influence of node degree, Jaccard index, HDI index and other similarity indicators based on local information are derived. Because of its high robustness and easy operation, these algorithms have attracted much attention of the researchers. In addition to extracting structural information, similarity index based on path information is also used in the link prediction. The similarity index based on path information includes Katz index and Local Path index (Lu, et al., 2009). Generally, when two proteins are predicted to be linked, they often share similar functions, suggesting that links are influenced by the interactions and the external neighbors. This concept represents a naive Bayesian model (Liu, et al., 2011). Additionally, node degree and the clustering coefficient impact on establishing of links by combining information from common neighbor nodes for prediction were also proposed (Yan, et al., 2016; Yang, et al., 2017).

## Acknowledgments

This work was supported by National Natural Science Foundation of China (61303108); The National Key Research and Development Program of China (Grant No. 2016YFC1306605), The Natural Science Foundation of Jiangsu Higher Education Institutions of China (17KJA520004); Suzhou Key Industries Technological Innovation-Prospective Applied Research Project (SYG201804); Program of the Provincial Key Laboratory for Computer Information Processing Technology (Soochow University) (KJS1524); A Project Funded by the Priority Academic Program Development of Jiangsu Higher Education Institutions.

